# Cis-regulatory divergence and mate recognition in behaviourally isolated cricket species

**DOI:** 10.1101/2025.05.19.654830

**Authors:** Leeban H. Yusuf, Shangzhe Zhang, Vivienne Litzke, Stefan R. Pulver, Theresa M. Jones, Nathan W. Bailey

**Affiliations:** Centre for Biological Diversity, School of Biology, University of St Andrews, St Andrews KY16 9TH, Scotland; School of Psychology and Neuroscience, University of St Andrews, St Andrews, United Kingdom; School of BioSciences, The University of Melbourne, Melbourne, Victoria 3010, Australia

**Keywords:** mate recognition, female preference, gene expression, sexual selection, speciation

## Abstract

Cis-regulatory evolution has been implicated in morphological and physiological reproductive isolation, but its role in divergence of mate recognition behaviours is unknown. The latter are often the first, most powerful mating barriers to evolve during speciation, but understanding their genetics is hampered by the context-dependence of mate choice. We circumvented this challenge by using allele-specific expression analyses in F_1_ hybrids of sexually isolated cricket species, *Teleogryllus oceanicus* and *T. commodus*. Female crickets choose mating partners based on species-specific male advertisement songs, and we found extensive cis-regulatory divergence in neural tissues exposed to conspecific, heterospecific, or mixed-species songs. Genes with divergent expression were implicated in learning, memory and transcriptional regulation, and they showed elevated population genetic differentiation in allopatry but not sympatry. This implies that genes recruited during evolutionary divergence of mate recognition systems may be homogenised by gene flow. Regulatory divergence was asymmetrical between the two species; divergent selection on genes associated with female perception of male song was more prominent in *T. oceanicus* than in *T. commodus*. Evolved differences in gene regulation are strongly associated with sexual signal perception in closely related species, supporting the idea that rapid cis-regulatory evolution of mate recognition can accelerate speciation.

## Introduction

Coordinated divergence of sexual signals and corresponding preferences can generate assortative mating and sexual isolation (Lande, 1981; Kirkpatrick, 1982; Shaw *et al*., 2024), and result in some of the most rapid cases of speciation known (Mendelson & Shaw, 2005; Wellenreuther & Sánchez-Guillén, 2016). Despite this, remarkably little is known about the genetic mechanisms that cause differences in female preference between species (Butlin & Ritchie, 1989; Ritchie & Butlin, 2024). Genomic insights from the few studies that test the molecular basis of mate recognition in diverging species indicate that sexual signals and preferences are underpinned by complex genetic architectures, are likely under strong divergent selection, and may also be under divergent ecological selection (Shaw & Lesnick, 2009; Bay *et al*., 2017; Blankers *et al*., 2019; Rossi *et al*., 2020, 2024; Unbehend *et al*., 2021). However, the genetic mode of action driving differences in signals and preferences between recently diverged species is unknown in all but a handful of cases (Combs *et al*., 2018; Unbehend *et al*., 2021; Rossi *et al*., 2024).

Gene expression evolution – in particular, cis-regulatory evolution – has been argued to underpin divergence in morphological and physiological reproductive isolation between nascent species (Mack & Nachman, 2017). However, divergence in mate recognition and choice is rarely attributed to cis-regulatory evolution. A technical challenge may account for this: typical gene expression studies do not distinguish plastic, context-dependent changes in gene expression from genetic, evolved differences in expression. For example, within species it is known that brief exposure to potential mates or mating signals can change the expression of genes associated with olfactory signalling, immunity, and synaptic activity (Lawniczak & Begun, 2004; Woolley & Doupe, 2008; Immonen & Ritchie, 2012; Bloch *et al*., 2018; Keagy *et al*., 2024). Such studies suggest that genes associated with the expression of female preference might be activated immediately in response to male acoustic signals, making gene expression experiments a practical tool to investigate evolutionary divergence of female preference (Lynch *et al*., 2012; Ramsey *et al*., 2012; Bloch *et al*., 2018; Fischer *et al*., 2021). However, the fundamental challenge of detecting regulatory evolution between species is in distinguishing transcriptional responses to sexual signals that differ due to evolved, genetic changes driven by divergent selection during speciation, from gene expression responses that differ for incidental or nuanced reasons, including variation in development, cell type abundances and tissue allometry between species (Combs *et al*., 2018; York *et al*., 2018; Fraser, 2022). To eliminate these confounds, it is necessary to compare each species’ transcriptional responses in an identical genetic, developmental, and environmental background.

We tested how neural gene expression of females exposed to male sexual advertisement song has evolutionarily diverged between the field cricket species pair *Teleogryllus oceanicus* and *T. commodus* by performing a parental allele-specific expression study in F_1_ hybrids. These sister species are a classic system in research on acoustic communication and mate recognition (Hoy, 1974; Hoy *et al*., 1977). They are distributed across northern and eastern Australia, where they overlap in a zone of sympatry (**Fig. 1A**), and males of both species produce a conspicuous calling song to attract conspecific females. Male song and female mating preferences have diverged between *T. oceanicus* and *T. commodus*, resulting in a pre-mating barrier to gene flow, though hybrids can be readily formed in the lab (**Fig. 1A**) (Bailey *et al*., 2017; Moran *et al*., 2018, 2019). Species-specific differences in male calling song appear to occur via changes in conserved cellular and neuronal networks (Jacob & Hedwig, 2019), but differences in female preference appear to be controlled by divergent filtering mechanisms in *T. commodus* and *T. oceanicus* (Bailey *et al*., 2017). Key components of male song in *Teleogryllus* have a polygenic architecture: different components of male calling song are genetically modular and some genetic modules associated with male song are under divergent selection (Yusuf *et al*., 2024). However, little is known about the genomic basis of female perception of male sexual signals, whether it differs between *T. oceanicus* and *T. commodus*, and whether gene expression responses associated with female choice differ based on acoustic context (Bailey, 2011; Pascoal *et al*., 2018).

**Fig 1.**
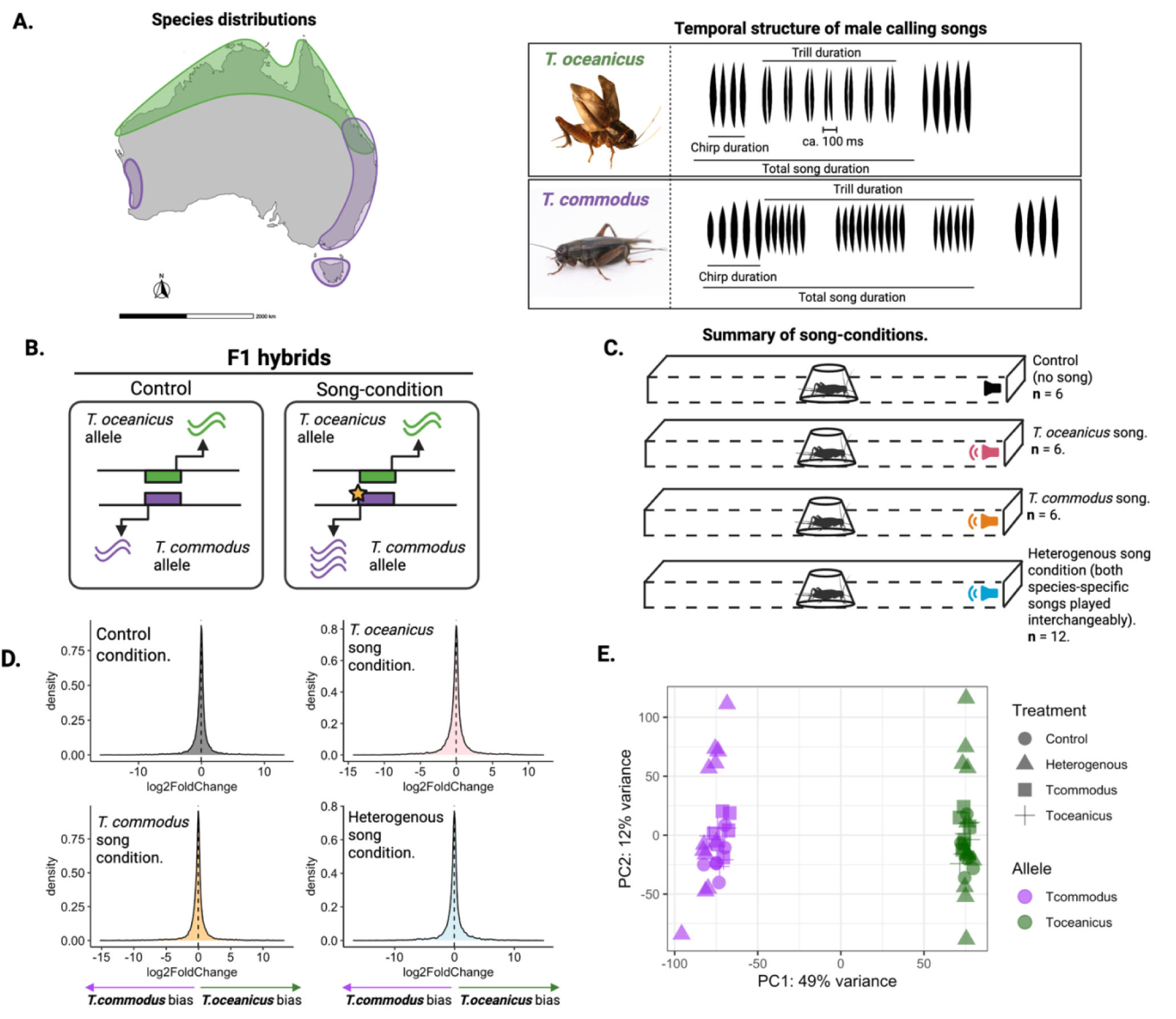
Description of experimental design. **A.** Australian distributions for *T. oceanicus* and *T. commodus* based on Otte & Alexander (1983) and Moran *et* (2018), with calling song diagrams based on the latter. Cricket photos: NW Bailey. **B.** Schematic showing potential cis-regulatory divergence related to acoustic signal perception can be detected using allele-specific ression assays in F1 hybrid females. Species-specific gene cis-regulatory regions are denoted by coloured angles. In the control, there is no difference in transcript abundance between *T. commodus* and *T. oceanicus*, when exposed to song, transcription differences between species can arise due to species-specific mutations ring transcription factor binding (denoted by the star on *T. commodus* regulatory region). **C.** Diagram of the erimental design. F1 hybrid females were either exposed to silence (control), species-specific song (*T. modus* or *T. oceanicus* song), or a heterogenous condition where both species-specific songs were alternated ng playback. Each F1 hybrid was exposed to one of these four conditions for 30 minutes. The number of ogical replicates is shown next to each condition (see also **Supplementary Table 1**). **D.** Observed ribution of allelic expression for all genes and conditions for F1 female hybrids. **E.** PCA of whole-brain scriptomics of all F1 hybrid female samples used in the analysis, categorised by species (purple = *T. modus* allele; green = *T. oceanicus* allele) and treatment.

Our first goal was to determine whether cis-regulatory evolution underlies female responses to sexual signals in diverged *Teleogryllus* species; and if so, whether this evolution affected shared or distinct gene expression networks in each (**Fig. 1B**). Genes expressed in an allele-specific manner between parental genomes contained within a single hybrid individual must be differentially expressed due to proximate (i.e. in cis-) sequence differences in one or both parental genomes, as environmental, developmental and cell-type abundance confounds are prevented. To elicit a transcriptional response, we exposed F_1_ hybrid females to male advertisement song for 30 minutes. We simulated allopatric and sympatric mate choice contexts by exposing females to either parental species’ advertisement song or to both species’ songs played in alternating sequence (**Fig. 1C**). Finding support for a mixture of shared and unique cis-regulatory evolution in each species, our next goal was to determine whether the implicated gene networks show greater differentiation and signatures of diversifying selection in wild populations. We generated a high-quality, annotated chromosome-level genome assembly for *T. commodus* and with an existing *T. oceanicus* reference genome (Zhang *et al*., 2024) analysed 60 whole-genome resequenced individuals from allopatric and sympatric populations in the wild. Patterns of genomic divergence and selection on genes implicated in cis-regulatory evolution indicate an important role in the diversification of female preferences and premating isolation, but with the caveat that cis-regulatory divergence is susceptible to degradation in sympatry due to gene flow.

## Results and Discussion

### Cis-regulatory divergence in neural transcriptomes is widespread and context-dependent

Our analyses identified allele-specific expression (ASE) differences – that is, differences in expression of gene copies from maternal vs. paternal genomes – within our control condition and song conditions (**Fig.1B**). The distribution of allele-specific expression was slightly biased towards *T. commodus* across all conditions (**Fig. 1C**). This is unlikely to be due to mapping biases since F1 hybrid individuals were produced using parents from the same laboratory populations as individuals used to assemble the genomes to which transcriptomic data were mapped. Specifically, we assembled a chromosome-level *T. commodus* genome assembly with a total length of 2.22 Gb and fourteen pseudo-chromosomes (**Supplementary Fig.1**), consistent with existing genome assemblies for *T. oceanicus* (Pascoal *et al*., 2020; Zhang *et al*., 2024). The *T. commodus* genome assembly includes a total of 1,107.80 Mb of transposable elements (TEs), which make up 51.87% of its total length. The most abundant TE class was DNA repeats, spanning 198.07 Mb and accounted for 17.63% of the annotated TEs in the *T. commodus* genome. The final assembly used in the following analyses had a scaffold N50 of 139.51 Mb and BUSCO completeness of 96.8% ([S:65.1%, D:31.7%], F:0.6%, M:2.6%). Our high-quality contiguous genome for *T. commodus* represents a significant important addition to burgeoning genomic resources for *Ensifera* and will be useful for future comparative analyses across singing insects.

We accounted for the global bias towards *T. commodus* in our design by removing any allele-specific genes also found to be differentially expressed between parental genomes in the control condition, thus minimising any expression differences arising from technical (e.g., mapping bias (**Supplementary Fig. 2**)) or biological factors (e.g., hybrid misexpression) not caused by a response to song stimuli. Across all conditions, F1 hybrid female neural samples grouped by parental allele, indicating strong cis-regulatory divergence in neural expression patterns between species, irrespective of acoustic environment (**Fig. 1D**).

By analysing neural ASE patterns, we discovered that cis-regulatory evolution between *T. oceanicus* and *T. commodus* generally affected nearly a quarter of genes in the genome. Overall, we analysed 25,331 genes, and in the control condition 5,964 of these (23.5%) showed ASE. After removing this control set to focus only on genes whose responsiveness to sexual advertisement song has diverged, we found: 864 ASE genes after exposure to *T. oceanicus* song (3.4%), 589 ASE genes after exposure to *T. commodus* song (2.3%) and 2,217 ASE genes after exposure to both species’ songs (8.7%). We found no strong evidence for allelic bias towards either parental genome when comparing single-species song conditions against the control (Fisher’s exact test: Control - *T. oceanicus* song, *p* = 0.63; Control – *T. commodus* song, *p =* 0.25), but in the heterogenous song condition we observed significantly more *T. commodus* biased genes than *T. oceanicus* biased genes compared to the control (Fisher’s exact test: Control – Heterogenous song, *p <* 0.001) (**Supplementary Fig. 3**). Thus, the scale of cis-regulatory divergence related to song perception between parental genomes is consistent across behavioural states, affecting between ca. 2 – 9% of genes in the genome (**Fig. 2A**).

**Fig. 2.**
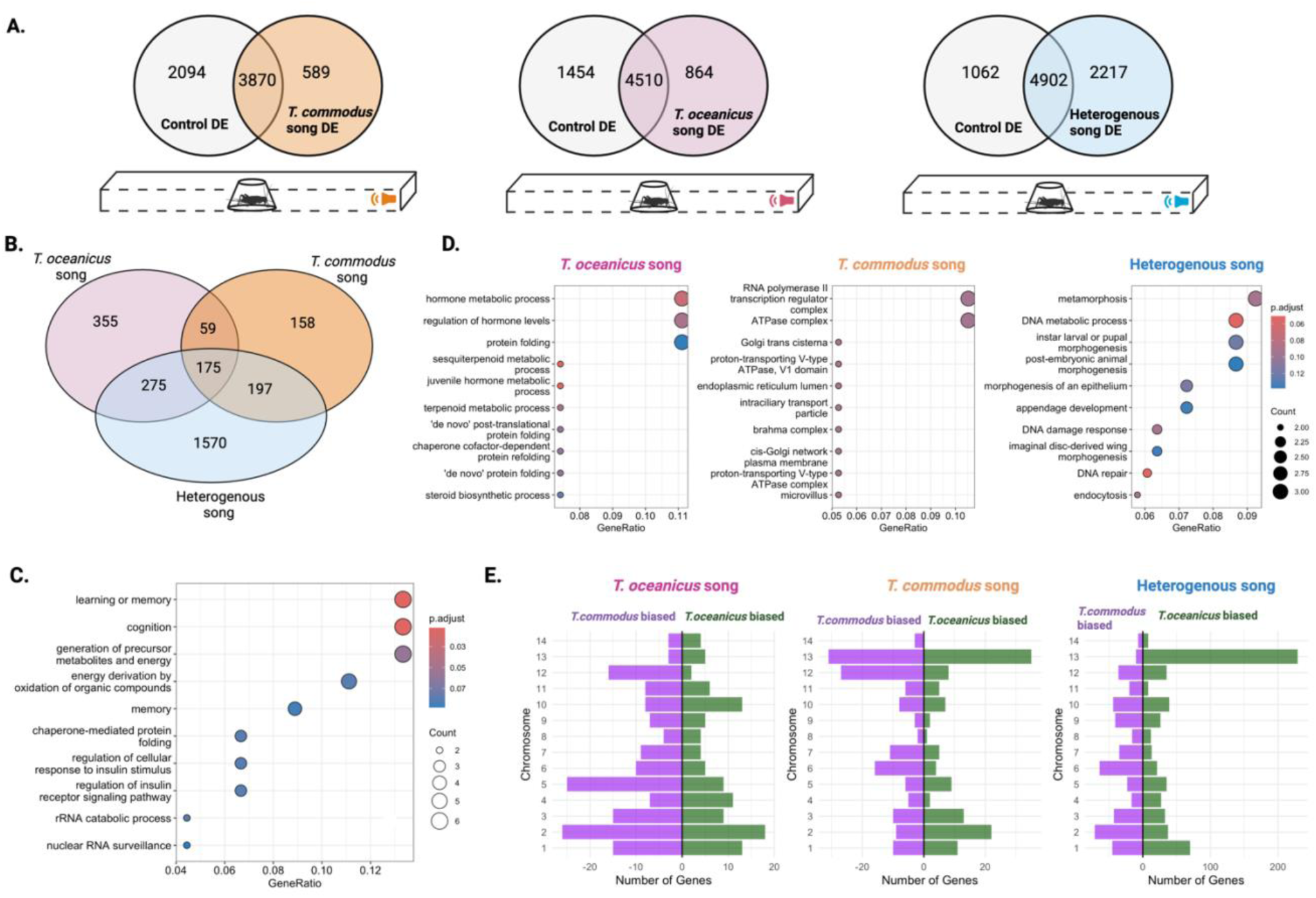
Allele-specific expression in female *Teleogryllus* hybrids in response to male song. **A.** Venn diagrams showing number of genes with significant ASE in pairwise control vs. song comparisons. Only those genes that were uniquely ASE in each treatment, but not in the control condition, were retained for further analysis. **B.** Venn diagram showing overlap between retained ASE genes in each song treatment. **C.** Biological GO terms for ASE genes shared across all song treatments. **D.** Gene ontology showing biological enrichments for uniquely expressed ASEs in each song treatment indicating that the latter have different functions in different treatments. **E.** Chromosomal distributions of ASE genes in each song condition.

### Cis-regulatory divergence across song conditions: Shared ASE genes

We next tested the extent to which ASEs were shared vs. unique across male song conditions. This analysis allowed us to distinguish whether responses to conspecific vs. heterospecific sexual signals activates a common set of genes with divergent expression between species – suggesting that cis-regulatory divergence governs a common response to sexual signal variants – or whether such responses utilise different gene sets that underwent cis-regulatory divergence. These are not mutually exclusive, but disentangling their relative importance enables inference about whether sexual isolation evolves via shared or discrete genetic pathways in diverging species, which has implications for understanding the consequences of gene flow, hybridisation, and the stability of species boundaries (Felsenstein, 1981). We predicted that genes showing ASE would differ in each song condition, because the decision rules governing female behavioural responses to conspecific versus heterospecific signals differ between *T. oceanicus* and *T. commodus* (Bailey *et al*., 2017). To evaluate this prediction, we tested whether more ASE genes were shared across song conditions than expected by chance (**Fig. 2B**). They were (hypergeometric test; n = 234; p < 0.001), which rejected the idea of wholly distinct genetic architectures undergoing separate cis-regulatory evolution in the two species.

A core transcriptional programme contributing to species recognition appears to have genetically diverged between *T. oceanicus* and *T. commodus*. We observed a significant overlap in ASE genes expressed when F1 hybrids respond to heterogenous song and *T. oceanicus* song conditions (n = 450; hypergeometric test, p < 0.001), and between heterogenous song and *T. commodus* song conditions (n = 372; hypergeometric test, p < 0.001) (**Fig. 2B**). Using gene ontology (GO) enrichment analyses, we tested whether the cohort of significant ASE genes shared across all song conditions (n = 175) were dominated by any particular biological functions. Only two GO categories related to biological processes were significant following multiple testing correction (**Fig. 2C**) (FDR < 0.05). These included genes involved in learning and memory (FDR = 0.01; p < 0.001), and cognition-related genes (FDR = 0.01; p < 0.001), indicating that cis-regulatory evolution affecting divergent preferences between species affects genes involved in long- or short-term neural processing of male song. Other enriched terms related to cellular processes include: oxidoreductive processes (FDR = 0.04; p < 0.001) and the nuclear exosome (FDR = 0.04; p < 0.001).

Six ASE genes were shared across all song conditions and implicated in learning and cognition. For example, *shaggy* (sgg) (*T. commodus* biased) is a glycogen synthase kinase that modulates olfactory habituation and circadian locomotor activity (Martinek *et al*., 2001; Yuan *et al*., 2005; Wolf *et al*., 2007). Similarly, the Wnt pathway co-receptor *arrow* (*T. commodus* biased) is essential for memory in *Drosophila*, where its inhibition impairs long-term memory (Wehrli *et al*., 2000; Tan *et al*., 2013). *Tomosyn* (*T. oceanicus* biased) regulates the release of neurotransmitters, hormones, proteins and micoRNAs, impacting memory and learning efficiency in associative tasks (Yizhar *et al*., 2004; Sakisaka *et al*., 2008; Yamamoto *et al*., 2010; Subkhangulova *et al*., 2023), and *krasavietz* (*T. commodus* biased)*, Ceng1a (T. oceanicus biased)* and *fdh* (*T. commodus* biased) influence behaviours related to motor function, synaptic plasticity and spatial learning (Lee *et al*., 2007; Acebes *et al*., 2012; Gross *et al*., 2015; Ai *et al*., 2019). A key similarity shared by most of these learning genes is that they are responsive to olfactory cues in *Drosophila*, corroborating previous evidence that genes involved in chemical communication may be co-opted for acoustic pattern recognition or have pleiotropic effects on acoustic signal processing (Immonen & Ritchie, 2012).

By examining divergent expression across all three conditions we have identified new candidate genes not previously linked to female preference or choice. Several genes previously linked to courtship song in *Drosophila* also show divergent expression between *T. commodus* and *T. oceanicus* in at least one song treatment (**Supplementary Table 2**). Namely, *fruitless* (*fru*) showed divergent expression between species during the *T. commodus* (log_2_FC =1.12; *p* = 0.04) and heterogenous (log_2_FC = 2.87; *p* < 0.001) song treatments. *fru* is a transcription factor that specifies sex-specific behaviours including male courtship (Rideout *et al*., 2007), and appears to affect female receptivity in *Drosophila* (Chowdhury *et al*., 2020). Similarly, the gene *period* (*per*) showed divergent expression in two song conditions, but not in the control treatment (*T. oceanicus* song: log_2_FC = 0.36, *p* = 0.02; heterogenous song: log_2_FC = 0.52*, p* = 0.04). Like *fru*, the gene *per* has long been known to affect male courtship song, as well as circadian rhythms (Kyriacou & Hall, 1980; Greenacre *et al*., 1993; Kyriacou, 2002). Our analysis demonstrates that genes with prominent effects on male and female courtship in *Drosophila* are potentially implicated in sexual isolation in other insects.

### Cis-regulatory divergence across song conditions: Unique ASE genes

A subset of ASE genes were unique to each song condition (**Fig. 2B**), representing cis-regulatory evolution of different components of neuronal processing or behavioural filtering, as well as context-dependent female responses. These condition-specific ASE genes diverged in function (**Fig. 2D**). In response to *T. oceanicus* song, species-specific expression differences related to hormone production (*p* = 0.07) and regulation (*p* = 0.09), including juvenile hormone regulation (*p* = 0.05). In response to *T. commodus* song, GO enrichment indicated divergent expression of the transcriptional machinery (*p* = 0.01), whilst in the heterogenous condition, we observed cis-regulatory expression divergence in DNA metabolic processes (*p* < 0.01) and metamorphosis (*p* < 0.01). Across all song conditions, divergent cis-regulatory genes responding to specific acoustic contexts have a polygenic genetic architecture (**Fig. 2E**). However, there was little consistency in the distribution of ASE genes by condition or by the direction of allelic bias (i.e. *T. commodus* biased or *T. oceanicus* biased) in expression patterns. This does not support the idea that cis-regulatory divergence of sexual signals and preferences are concentrated in particular ‘genomic’ hotspots.

### Cis-regulatory divergence vs. genomic divergence at song recognition loci

Divergence in cis-regulatory expression of genes associated with female response to male song would be consistent with a role for these genes during species divergence. We tested this idea by asking whether ASE genes show corresponding patterns of divergent genomic selection using population genomic data for *T. oceanicus* and *T. commodus*. We re-sequenced 60 *T. commodus* and *T. oceanicus* individuals (58 females and 2 males) from allopatric and sympatric populations on the eastern coast of Australia, where they overlap in a contact zone (**Fig. 1A; Supplementary Table 6**). The species are well-separated genetically and show no evidence of hybrid individuals in the wild (**Fig. 3A**). Then we compared multiple population genomic measures to evaluate signatures of divergent selection (genetic differentiation (F_ST_), Tajima’s *D* and local admixture), testing whether all ASE genes show enhanced divergence by comparing their patterns of genetic variation to background, genome-wide expectations.

**Fig 3.**
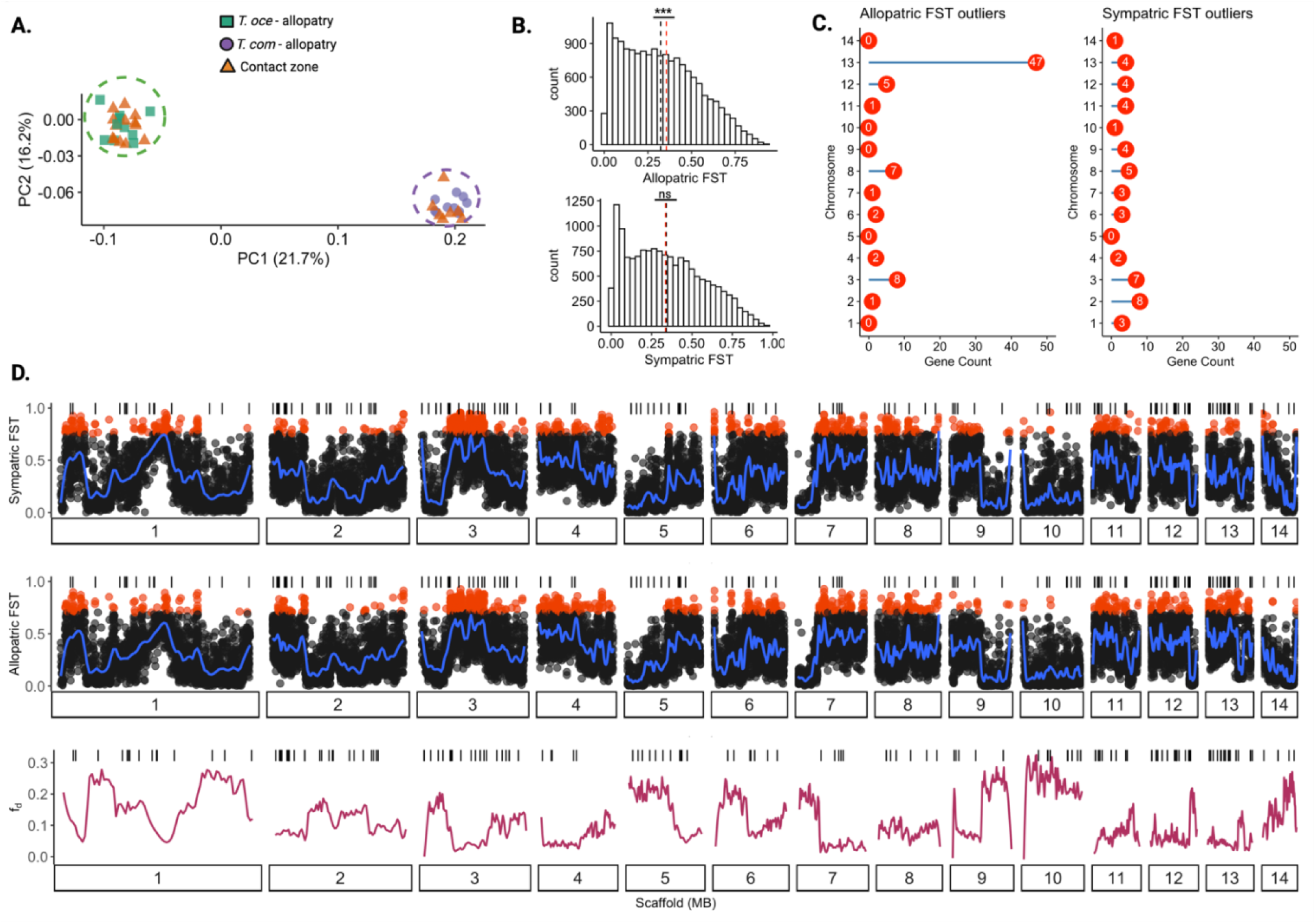
Genome-wide patterns of genetic differentiation associated with ASE genes in females. **A.** PCA based on whole-genome resequencing data showing genetic distance between *T. oceanicus* and *T. commodus* sampled from allopatry and sympatry. **B.** Histogram of F_ST_ calculated in 10kb windows among allopatric individuals only (top) and sympatric individuals only (bottom). In each comparison, the black dotted line illustrates the genome-wide mean and the red dotted line illustrates mean F_ST_ for all ASE genes across all song conditions. **C.** The number of ASE genes within F_ST_ outlier regions (red dots – 95^th^ quartile) by chromosome in both allopatric and sympatric comparison. **D.** Genome-wide F_ST_ for allopatric and sympatric comparisons, and genome-wide local admixture (f_d_) between *T. oceanicus* and *T. commodus*. Vertical lines above each genome-wide plot illustrate female “response” genes that showed allelic divergence across all song conditions (n = 175)

ASE genes associated with female responses to male song showed strong genomic divergence, but this was subject to genetic homogenisation in sympatry by occasional gene flow. In allopatry, ASE genes show significantly higher F_ST_ compared to the genomic background (one-sample Wilcoxon test; *p* < 0.001), but in sympatry this difference disappears (one-sample Wilcoxon test; *p* = 0.08) (**Fig. 3B**). Using a measure of local admixture (f_d_) we compared levels of gene flow between all *T. oceanicus* and *T. commodus* individuals in our population genomic dataset. ASE genes show lower average levels of gene flow (f_d_ = 0.12) compared to the rest of the genome (f_d_ = 0.14), which is consistent with divergent selection acting on genes involved in female response to male song (one-sample Wilcoxon test; *p* < 0.001).

To further explore this pattern, we compared F_ST_ between *T. oceanicus* and *T. commodus* females for ASE genes biased toward *T. commodus* vs. those biased towards *T. oceanicus*, for each song treatment. A difference in genomic differentiation (F_ST_) between *T. commodus*-biased vs. *T. oceanicus-*biased ASE genes, and in specific song treatments, would strongly imply that (*i*) ASE genes may be driven by directional selection in one particular species, and (*ii*) that female responses in particular mate choice scenarios may be more relevant to sexual isolation than others. In allopatry, *T. oceanicus*-biased ASE genes showed significantly higher genetic differentiation than *T. commodus*-biased genes for ASE genes identified in the heterogeneous and *T. commodus* song conditions, but not in the *T. oceanicus* song condition (Wilcoxon test: *T. oceanicus* song, *p* = 0.43; *T. commodus* song, *p* < 0.01; heterog**e**neous song, *p* < 0.01). In sympatry, however, we found little support for differentiation between *T. commodus*-biased vs. *T. oceanicus*-biased ASE genes across song conditions. These results imply, as above, that any signal of divergent selection acting on cis-regulatory divergence seems to be specific to allopatry, cis-regulatory divergence involved in female perception of male song may be homogenised by gene flow in sympatry, and that divergent selection may be context-specific.

To corroborate nuanced patterns of selection, we next asked whether ASE genes showed species-specific signals of positive selection by comparing Tajima’s *D* in a species-specific manner. Given differences observed between allopatric populations but not sympatric populations, we compared Tajima’s *D* in allopatric populations of *T. commodus* and *T. oceanicus* to background genomic levels of Tajima’s *D* in each species, expecting significantly lower values in ASE genes if they have been subject to recent positive selection. Tajima’s *D* was no different from the genome-wide mean in *T. oceanicus* ASE genes (one-sample Wilcoxon test; *p* = 0.93), and was significantly higher in *T. commodus* ASE genes (one-sample Wilcoxon test; *p* < 0.001). Overall, our results are consistent with a scenario of older (sexual) selection driving patterns of genomic differentiation in ASE genes given little evidence of recent, positive selection on ASE genes on average and a weakening of sexual barriers in sympatry.

### Potential barrier loci implicated in female response to male advertisement song

We next asked how many of all ASE genes identified across our song conditions were in outlier regions characterised by high levels (95^th^ quartile) of F_ST_, consistent with a potential role in speciation. In allopatry, 74 ASE genes were found in regions of high genetic differentiation compared to 49 genes in sympatry (**Fig. 3C**). We also found a significant positive correlation between the number of ASE genes per chromosome and mean genetic differentiation per chromosome in allopatry (*r* = 0.74, *p* < 0.01), but not in sympatry (*r* = 0.34, *p* = 0.24). These results suggest that a genome-wide reduction of genetic differentiation in sympatry – likely the result of occasional gene flow between *T. oceanicus* and *T. commodus* – also has a homogenising effect on genetic variation linked to female response genes. This effect is most pronounced on chromosome 13, where we observe 47 out of 74 ASE genes in outlier windows of genetic differentiation in allopatry, but only four of those genes remain as outliers in sympatry (**Fig. 3C**).

A shared, core set of ASE genes (N = 175) were shared across all song conditions and may be expected to reflect context-independent neural or behavioural response pathways. As with the full cohort of genes showing significant allele-specific expression in each treatment, this smaller shared subset of female ‘response’ genes are distributed genome-wide (**Fig 3D**). Four genes in this shared core gene set were also outliers in a genome scan of genetic differentiation between the two species in both allopatry (QTRTD1, *ben*, *EbpIII* and *Tango*5) and sympatry (*Tango5* and CIAPIN1, two unannotated). Of these, *ben* encodes a ubiquitin-conjugating enzyme and is well-known to be involved in synapse formation and synaptic growth; mutations in *ben* are linked to contextual learning deficits and changes in locomotion in *Drosophila* (Muralidhar & Thomas, 1993; Oh *et al*., 1994; Uthaman *et al*., 2008). Another gene of interest is *EbpIII* – a pherokine typically expressed in olfactory tissues. *EbpIII* has been implicated in chemical communication in insects, forms part of the posterior mating plug which influences female remating rates in *Drosophila*, and mediates parent-offspring interactions in earwigs (Lung & Wolfner, 2001; Sabatier *et al*., 2003; Bretman *et al*., 2010; Wu *et al*., 2020).

There are only a few detailed examples of candidate genes involved in mediating preference for chemosensory, visual or auditory cues across animals. One particularly interesting candidate gene – *regucalcin1* – satisfied the following stringent conditions: it showed pronounced genetic differentiation between *T. commodus* and *T. oceanicus* in allopatry (F_ST_ = 0.80) and sympatry (F_ST_ = 0.82); there was parental allele-specific expression in response to species-specific song (*T. commodus* song: *p* < 0.01; *T. commodus* song: *p* < 0.01) but not heterogenous song (**Supplementary Fig. 4**). Local levels of gene flow between *T. commodus* and *T. oceanicus* at *regucalcin1* are markedly depressed compared to background genomic levels on Scaffold 3 (**Supplementary Fig. 5A**). Previous analyses have demonstrated that allele-specific expression and population genomic divergence of *regucalcin1* is strongly associated with visual mate preference in *Heliconius*, and disruption of *regucalcin1* affects courtship (Rossi *et al*., 2020, 2024). We detected evidence of a strong selective sweep in *regucalcin1* in *T. oceanicus*, but not in *T. commodus*, implying that the observed cis-regulatory divergence of *regucalcin1* is specific to *T. oceanicus* (**Supplementary Fig. 5B**). Our analyses suggest that convergent molecular evolution of neural processing/integration genes such as *regucalcin1*, alongside genes involved in detection of sexual signals, could provide a common mechanism for divergence of sexual communication between new species.

## Conclusion

The extent to which gene expression differences between species contribute to divergence of mate recognition and discrimination – often the first barrier to evolve between nascent species (Shaw *et al*., 2024) – has been largely unknown. We found extensive, genome-wide cis-regulatory expression divergence associated with responses to male acoustic sexual signals in reproductively isolated field cricket sister species of the genus *Teleogryllus*. This cis-regulatory divergence underpins responses in a realistic sympatric context where females are exposed to both species’ sexual signals at once. The latter is important, because where these species overlap in sympatry, they show no evidence of niche differentiation and often sing side-by-side simultaneously (Moran et al. 2020). There is strikingly more cis-regulatory gene expression divergence associated with this complex, simultaneous-choice situation than in either single-species signalling context, implying different neurogenetic programmes of mate recognition in different social contexts. Cis-regulatory divergence underlying sexual isolation via premating mate choice is supported by multiple lines of evidence including the broad distribution of diverged cis-regulatory genes across chromosomes, differences in the functions of genes with divergent expression in different contexts, and genomic signatures of divergent selection in allopatry. Our findings demonstrate that regulatory evolution can explain dynamic and constitutive gene expression changes underlying differences in complex mate choice decisions between species.

## Methods

### Animal husbandry

We obtained *T. commodus* males from stock populations originally collected from Bluey’s Beach, New South Wales, Australia, and normal-wing/wild-type *T. oceanicus* females from Kauai, Hawaii, USA. We crossed 10 *T. commodus* males and 10 *T. oceanicus* females in 10 independent crosses to produce multiple replicate F1 populations and randomly selected one F1 hybrid population for experiments. The reciprocal cross was not performed because this direction suffers from lower offspring numbers (because of probable partial sterility/inviability) based on previous analyses (Moran *et al*., 2017). Individuals from the selected F1 hybrid population were housed in a plastic storage container until their sex was identifiable at their final instar. During their final instar, female hybrids were isolated from males and kept in separate 113 mL deli pots and were maintained biweekly with food, an egg carton and a small glass-water vial. From isolation at their final instar, F1 hybrid females were monitored every other day to check for eclosion and were housed away from males in a sound-proofed incubator to prevent them from hearing any male advertisement and courtship songs. Incubators were set at 25 °C on a photo-reversed 12:12 light/dark cycle.

### Experimental song trials

To understand interspecies gene expression differences in female response to male song, we assayed gene expression in hybrid female brains following exposure to three male calling treatments. Specifically, we exposed F1 hybrid females to one of three male advertisement song conditions 7-10 days post-eclosion via song playback. Six individuals were exposed to silence (our control), six individuals were exposed to *T. oceanicus* song, six individuals were exposed to *T. commodus* song, and finally, 12 hybrid females were exposed to song containing each species-song interchangeably. Within each treatment, females were exposed to two different species-specific song, so that three individuals were exposed to one intraspecific song and the other to another intraspecific both from the same species. Both songs were population-specific synthetic models with slightly different parameters. This was to ensure that we could distinguish between allelic expression as a response to species-specific song rather than any specific features of a given song belonging to an individual. Two population-specific synthetic song models for *T. commodus* were taken from Bailey & Macleod, (2014).

Briefly, song models were constructed by recording male calling songs using a Sennheiser ME66 microphone under red light between 22-25°C (with one individual recorded at 19°C). For each individual, 10 song phrases from a continuous recording of male calling song were manually analysed using Sony SoundForge (v.7.0a). Parameters for song models were averaged between two populations from New South Wales and subsequently adjusted to match four key advertisement song parameters for the Blueys Beach stock population (“number of chirp pulses” = 7, “number of pulses in trill 1” = 15, “number of pulses in trill 2” = 10, and “number of pulses in trill 3” = 8). For *T. oceanicus,* population-specific song models were constructed by averaging song parameters from two source populations from Daintree in Queensland, Australia, and Wailua in Kauai, Hawaii. We reasoned that *T. oceanicus* song models should reflect the fact that stock populations from Kauai, Hawaii may show different preferences for song compared to the source population that they were originally collected from after generations of inbreeding.

Each F1 hybrid female was placed and kept stationary in a small plastic pot in a designated position in a rectangular testing arena (275cm, 40cm, 28cm) during the course of each playback. All the playback experiments were performed under red light, at between 23-27°C in a soundproofed room. Between each playback experiment, the entire arena, plastic cups and all equipment were wiped down with 70% EtOH to reduce the likelihood of olfactory cues being transferred between F1 hybrid females. For each playback and for every individual, we used the same speaker (Logitech Z120) and the same MP3 player (SanDisk SDMX26-008G-E46P). The playback experiment for all conditions (except the control) were structured in the same way: females were permitted 1 minute to acclimate and then the allocated song was played for 5 minutes, followed by a 1-minute silent interval. We repeated this for a total of 30min (5 song repeats) after which time the female was immediately euthanised using sterilised dissecting scissors and the head transferred to screw-cap Eppendorf tubes containing RNAlater. All heads were placed into a -20°C freezer for storage prior to extraction.

### RNA extraction and sequencing

Female F1 hybrid heads were processed with one of two homogenisation approaches: A) heads were transferred into a 1.5 mL Eppendorf with of 300 µL TRIzol and homogenised via a hand-held homogenizer (Cole-Parmer® Motorized Pestle Mixer) for 5 minutes and left to incubate at room temperature for 5 minutes. Alternatively, B) heads were put into a 1.5 mL RNAse-free RINO® screw-cap tube and homogenised 12 for 3 minutes using a Bullet Blender® X24 and incubated at room temperature for 5 minutes. Following homogenisation, we extracted total RNA using TRIzol (Invitrogen) and Qiagen RNAeasy kit. All samples were stored in -80°C. mRNA libraries were prepared using PolyA selection and sequenced on an Illumina NovaSeq 6000 to produce paired 150bp reads. Library preparation and sequencing was performed at Novogene.

### DNA extraction and sequencing

Individual crickets were collected in 2013 from Australian populations detailed in Moran *et al*., (2018). Here, we sequenced 10 individuals from an allopatric population of *T. oceanicus* (Townsville, QLD) and an allopatric population of *T. commodus* (Bluey’s Beach, NSW), and individuals from a contact zone population containing both species (Tannum Sands, QLD). In total, we include 40 individuals for population genomic analyses in this study. Genomic DNA was extracted from single hind legs using the CTAB method. The quality and quantity of extracted DNA was measured using Qubit dsDNA kit and Nanodrop. Library preparation and sequencing was performed at Novogene, and sequencing was performed on an Illumina NovaSeq 6000 to produce paired end 150bp reads.

### Genome assembly and annotation

#### Sampling and DNA extraction

A female *T. commodus* individual was selected from a laboratory line originally collected from the wild in Australia (Blueys Beach, New South Wales).Cricket legs flash-frozen in liquid nitrogen were used for DNA extraction. First, the tissue was washed in 1% PBS buffer, and high–molecular-weight DNA was extracted using the Macherey-Nagel™ NucleoBond™ High Molecular Weight DNA Kit. The quality and quantity of the extracted DNA were assessed using both a Nanodrop™ spectrophotometer and a Qubit™ fluorometer, and the fragment length distribution was evaluated by electrophoresis. *De novo* genomic sequencing was performed by the Beijing Genome Institute (BGI) using the PacBio Sequel^®^ platform. In total, 7,951,018 HiFi reads were generated, yielding approximately 103 Gb of sequence data, which covered the haplotype genome to a depth of about 50×. Using the same extraction method, a DNBseq library was constructed and sequenced on the MGI DNBSEQ™-T7 platform at BGI, producing c.a. 120 Gb of clean paired-end reads. Additionally, cricket legs flash-frozen in liquid nitrogen was used for Hi-C library preparation and sequencing at BGI, generating approximately 120 Gb of Hi-C raw reads.

#### Draft genome assembling

HiFiasm v0.19.5-r587 (Cheng *et al*., 2022) was used with default settings to generate a draft *T. commodus* assembly. The resulting assembly had a total length of 2.22 Gb, comprised 1,184 contigs, and had a contig N50 of 5.39 Mb. To remove potentially duplicated contigs, Purge_Dups v1.2.6 (Guan *et al*., 2020) was employed, aligning k-mers from the HiFi raw reads back to the assembly to identify and mark duplicate sequences. Mitochondrial sequences were identified using MitoHiFi v3.2.1 (Uliano-Silva *et al*., 2023). The complete mitochondrial genome of *Teleogryllus emma* (YE *et al*., 2008), downloaded from the NCBI RefSeq database, was used as the reference. In total, 26 contigs in the draft assembly were identified as mitochondrial sequences and removed. The longest circular contig was designated as the *T. commodus* mitochondrial genome and annotated using the GeSeq module of CHLOROBOX (Tillich *et al*., 2017). After removing the mitochondrial sequences, the final contig-level assembly was 2.13 Gb in size, with a contig N50 of 5.61 Mb.

#### Hi-C scaffolding

Following the Arima Hi-C mapping guidelines (https://github.com/ArimaGenomics/mapping_pipeline), all Hi-C reads were mapped to the contig-level assembly. YahS (Zhou *et al*., 2023) was then used to scaffold the contigs based on these mapping results. After the initial scaffolding, Juicer Tools (https://github.com/aidenlab/juicer) and Juicebox (https://github.com/aidenlab/Juicebox) were employed to visualise the contact heatmap and manually correct the scaffolding. The final scaffold-level assembly had a total length of 2.14 Gb. Fourteen pseudo-chromosomes were created, while 212 contigs remained unscaffolded, resulting in 226 total scaffolds with an N50 of 139.51 Mb. This assembly was used for all subsequent analyses.

#### Genome quality assessment

All BUSCO assessments were performed using BUSCO v5.8.2 (Manni *et al*., 2021) with the Insecta core gene set (insecta_db10). The “–genome” mode was used for the genomic sequences, and the “–transcriptome” mode was applied to the protein-coding gene set .In addition to the BUSCO completeness evaluations, two additional methods were employed to assess genome quality. First, Merquery v1.3 (Rhie *et al*., 2020) was used to generate k-mer spectra for both the HiFi reads and the assembly, providing an estimate of base accuracy and completeness. This analysis yielded a completeness of 91.20%, a QV of 59.45, and a base error rate of 1.13×10^-6^. Second, the mappability of the genome was tested by aligning HiFi reads with NGMLR(Sedlazeck *et al*., 2018) and DNBseq short reads with BWA-MEM2 (Vasimuddin *et al*., 2019). The mapping rates were 99.65% for HiFi reads and 99.30% for short reads, indicating that the genome is well-suited for subsequent analyses.

#### Genome annotation

To identify repetitive elements in the genome, a three-step approach was employed. First, Tandem Repeats Finder (TRF) v4.09.1 (Benson, 1999) was used to detect simple repeats, which were then hard-masked with ‘N’ in the genome sequence. Second, transposable elements (TEs) were identified using RepeatMasker v4.1.0 (Tarailo-Graovac & Chen, 2009) with the masked genome as input, aligning the sequences to Repbase (Bao *et al*., 2015) to detect Arthropoda-homologous TEs. The genome was subsequently hard-masked based on these TE identifications. Finally, RepeatModeler v2.0.5 (Flynn *et al*., 2020) was applied to align the genome against itself and identify additional repeats. These results were incorporated into a custom database, and RepeatMasker was run again to annotate repeats based on alignment hits. All repetitive elements identified by this procedure were soft-masked for subsequent analyses.

For gene prediction, we used both homologous protein sequences and RNA-seq data. Protein sequences from *Drosophila melanogaster* (BDGP6.46, Ensembl; Harrison *et al*., 2024) *Gryllus bimaculatus* (Ylla *et al*., 2021) *Teleogryllus occipitalis* (Kataoka *et al*., 2020) and *T. oceanicus* (v2, Zhang *et al*., 2024), and orthologs from the arthropoda_10 dataset were obtained from public databases. Additionally, RNA-seq data from previous research were retrieved from the European Nucleotide Archive (ENA). Prior to gene prediction, raw RNA-seq reads were trimmed with fastp v0.23.0 and aligned to the reference genome using HISAT2 v 2.2.1 (Kim *et al*., 2019).

We then applied a three-step, integrated prediction workflow: 1) BRAKER v3.0 was used to generate *ab initio* gene predictions by training gene models based on both protein and transcript evidence. 2) Protein sequences were mapped to the reference genome to predict potential gene coordinates and ORFs. GeMoMa v1.9 was employed for proteins from closely related species, while exonerate v2.4.0 was used to align Arthropoda core orthologs. 3) Transcripts were assembled with StringTie v2.2.0 and Trinity v2.15.0, respectively, both of which used the mapped RNA-seq data as input. For the Trinity-assembled transcriptome, sequences shorter than 1 kb were discarded, and the remaining transcripts were clustered with CD-HIT-EST v4.8.1 to remove redundancies. The PASA pipeline (https://github.com/PASApipeline/PASApipeline) was then used to align these transcripts to the reference genome and predict gene structures. For the StringTie-assembled transcriptome, potential ORF locations were predicted by TransDecoder v5.7.1 (https://github.com/TransDecoder/TransDecoder), and the resulting coordinates were converted to the reference genome. Finally, we integrated all gene predictions using EvidenceModeler v2.1.0 and refined the transcript structures with the PASA pipeline, obtaining the final gene set. The predicted gene number for the *T. commodus* genome was 23,375 and number of protein sequences including translation result of all transcripts is 35,333.

#### Functional annotation

The functions of protein-coding genes were annotated by mapping the isoform sequences against multiple publicly available functional databases. For Non-redundant (Nr; Sayers *et al*., 2021) and Swiss-Prot/TrEMBL (Boeckmann *et al*., 2003), protein sequences were mapped to an Insecta-specific subset using DIAMOND blastp v2.0.14.152 (Buchfink *et al*., 2021), enabling accurate identification of protein functions within this taxonomic group. To identify and annotate functional domains within the protein sequences, InterProScan (Paysan-Lafosse *et al*., 2023) was employed, which integrates multiple signature databases for a thorough domain analysis. Additionally, EggNOG mapper (Cantalapiedra *et al*., 2021) was utilised to conduct EggNOG (Huerta-Cepas *et al*., 2019) analysis, from which Gene Ontology (GO) terms were extracted to provide insights into the biological processes, molecular functions, and cellular components associated with each protein. Finally, KEGG pathway (Kanehisa *et al*., 2016) annotations were acquired using the KAAS online server (Moriya *et al*., 2007), allowing for the identification of biochemical pathways and the functional context of the proteins within broader metabolic networks.

### Allele-specific expression quantification

Raw reads were cleaned for quality control using fastp with default parameters. Following quality control and filtering, we mapped sequences for each individual to both species reference genomes (*T. oceanicus* and *T. commodus*) using STAR with default parameters (Dobin *et al*., 2013). We used a competitive read-mapping approach called CompMap originally described in Sánchez-Ramírez *et al*., (2021) to obtain allele-specific read counts in F1 hybrids (Sánchez-Ramírez & Cutter, 2021). Unlike other approaches to categorise allele-specific expression that identify useable SNPs and mask homozygous regions, CompMap sorts and counts reads at a given homologous transcript by comparing read-mapping statistics (alignment score and number of mismatches) using each parental alignment. Reads that are best aligned to one or the other parental genome are assigned to that parental reference genome. Reads that ambiguously map (i.e. reads that map equally well to both parental genomes) are also counted and distributed proportionally relative to the number of reads already assigned to each parental genome. Read mapping bias was assessed by calculating percentage of uniquely mapped transcripts against each parental genome using STAR. We found a slight bias of uniquely mapped reads towards *T. commodus* (65.9% uniquely mapped reads) relative to the *T. oceanicus* genome (59.4%) (**Supplementary Figure 2**). These differences are unlikely to be caused by differences in genetic distance since F1 hybrid RNAseq here is generated from the same strains as data used to perform genome assembly. However, below we detail how our design accounts for this.

To ensure that transcripts were comparable and sorted in the same order for each parental genome prior to read counting, we used Liftoff, an annotation mapping tool, to ‘lift’ transcript features from the *T. commodus* genome annotation to the *T. oceanicus* genome using default parameters and by including ‘-exclude_partial’ to remove lifted features that only partially mapped or had low sequence identity (Shumate & Salzberg, 2021). Subsequently, to guarantee that only transcripts found in both genomes were used, we intersected BED files containing transcripts from *T. commodus* and transcripts lifted over to *T. oceanicus* using ‘bedtools intersect’(Quinlan & Hall, 2010). The resulting intersected BED file was used to remove any transcripts uniquely found in one parental genome compared to the other. BED files for both parental genomes were then sorted according to the name of the transcript identifier; this was necessary so that counting via CompMap could be performed on homologous transcripts. Before any analysis of counts, we summed allelic counts for transcripts belonging to the same gene to yield a single count for each allele per gene. Allele-specific counts were then estimated using CompMap for each sample.

### Allele-specific differential expression analyses

We analysed differential expression in allele-specific counts across treatments using DESEQ2 (Love *et al*., 2014). For interpretability, we opted to analyse each condition separately. We subsetted count data to include samples from the same condition and then quantified differential expression between parental alleles (∼ Allele) for control, *T. oceanicus* song, *T. commodus* song and heterogenous song separately. For each condition, we calculated library size factors and filtered out genes which did not meet our criteria of having a library-size scaled count at least equal to 10 across 3 samples. We visualised expression distances between allele-specific counts across samples using a PCA and confirmed that across all conditions samples grouped by allele. For all comparisons, we used the negative binomial generalized linear regression model fitting and Wald test to identify differentially expressed genes. We applied shrinkage of log_2_ fold change estimates to obtain more accurate fold change estimates using the package *apeglm*, as implemented in DESEQ2 (Zhu *et al*., 2019). Significant differentially expressed genes were first identified using FDR-adjusted p-values at a cut-off of 5%.

To identify significantly differentially expressed genes in the song conditions, we filtered all genes that were significantly differentially expressed in the control condition and in the song conditions in a pairwise manner using *dplyr.* Importantly, we reasoned that this step would remove genes showing (a) general neural expression divergence between parental genomes in hybrids, but also (b) genes that may be DE because of read mapping biases since all samples and conditions should suffer from this, including control samples. Finally, we restricted our analysis to focus on significantly expressed genes in song conditions that showed a log_2_ fold change larger than 1 or below -1.

### Gene ontology analysis

Annotations from *T. commodus* genome were imported into R to perform gene ontology analyses. First, to increase power to detect relevant gene ontology categories (involved in sound perception, mate choice, etc.), we converted annotations into *D. melanogaster* homologs where a 1-to-1 orthology was identified using the package *babelgene* (Dolgalev, 2022). We performed gene ontology enrichment analysis using clusterProfiler on genes significantly differentially expressed between parental alleles and uniquely expressed in each song condition, and that are expressed in all song conditions (Yu *et al*., 2011). To do this, we used a *D. melanogaster* gene set as background and filtered ontologies using FDR-adjusted p-value cut-off of 5%.

### Population genomics of allele-specific expression divergence

Genes with allele-specific expression show evidence of cis-regulatory divergence between species, but whether these genes show population genomic signatures of pronounced genetic divergence or adaptive evolution is an open question. We explored whether genes with divergent gene expression between species showed evidence of positive selection in the wild. To test this, we generated whole-genome resequencing data for 60 individuals from populations spanning the range of both species on the eastern coast of Australia, including a sympatric population (**Supplementary Table X**). Raw reads were trimmed using *fastp* (Chen *et al*., 2018). Subsequently, cleaned reads were mapped, sorted, filtered for duplicated using *sambamba* and *picard,* and variant calling was performed using *bcftools* (Tarasov *et al*., 2015; “Picard toolkit,” 2018; Danecek *et al*., 2021). We filtered for minimum depth (10), minimum mapping quality (30) and maximum allowed missingness at a given site (0.5 or genotypes for 20 individuals) performed using *bcftools*. We then computed F_ST_ in 50kb windows with 10kb step sizes using *pixy* to identify regions of the genome that are strongly diverged between both species (Korunes & Samuk, 2021). Additionally, we computed Tajima’s *D* in 50kb windows using *VCFtools* to identify the selective regime of genes showing expression divergence identified in each condition (Danecek *et al*., 2011). In the case of each summary statistic, we set a cut-off to identify outlier genomic regions by determining the 95^th^ quantile.

Additionally, we used the *Dinvestigate* function in Dsuite to quantify local admixture (f_d_) between *T. oceanicus* (P2) and *T. commodus* (P3) populations in 100bp windows with step sizes of 100bp closely following bioinformatic protocol in Martin *et al*., (2015) (Malinsky *et al*., 2021). For the outgroup, we used a publicly available sequencing dataset of a *T. occipitalis* individual (Kataoka *et al*., 2020), and for the ingroup we used Hawaiian individuals of *T. oceanicus* (P1) that we included from a previous study (Kataoka *et al*., 2020; Zhang *et al*., 2021). We used local patterns of admixture to test whether genes showing allele-specificity in song conditions are, on average, experiencing less gene flow compared to the background genome.

## Acknowledgements

We would like to thank Renjie Zhang and Ana Drago for helping with rearing crickets. L.H.Y was supported by grants from the Natural Environmental Research Council for funding (NE/W001616/1 and a NERC Discipline Hopping Grant). Additionally, the work was supported by the Australian Research Council Discovery award DP210101915, awarded to T.M.J and N.W.B

## Author contributions

Conceptualisation: L.H.Y., N.W.B.; Methodology: L.H.Y., S.Z., V.L.; Analysis: L.H.Y, S.Z. ; Visualisation: L.H.Y, S.Z.; Original draft: L.H.Y with input from S.Z. and N.W.B.; Editing: all authors; Funding Acquisition: L.H.Y., S.R.P., T.M.J., N.W.B.

## Data availability

Transcriptomic hybrid data and population genomic data will be deposited in sequencing read archive in NCBI under project name: PRXXXXXXX. Genome assembly and genomic data used to assemble the *T. commodus* genome used will be deposited in XX. Scripts used for analysis can be accessed in https://github.com/LeebanY/CricketFemaleResponseProject. Datasets detailing genes with significant allele-specific expression and F_ST_ outliers can be found in **Supplementary Tables 3-9**.

## References

Acebes, A., Devaud, J.-M., Arnés, M. & Ferrús, A. (2012) Central adaptation to odorants depends on PI3K levels in local interneurons of the antennal lobe. Journal of Neuroscience, 32, 417–422.

Ai, L., Tan, T., Tang, Y., Yang, J., Cui, D., Wang, R., et al. (2019) Endogenous formaldehyde is a memory-related molecule in mice and humans. Communications biology, 2, 446.

Bailey, N. & Macleod, E. (2014) Socially flexible female choice and premating isolation in field crickets (Teleogryllus spp.). Journal of Evolutionary Biology, 27, 170–180.

Bailey, N.W. (2011) Mate choice plasticity in the field cricket Teleogryllus oceanicus: effects of social experience in multiple modalities. Behavioral Ecology and Sociobiology, 65, 2269– 2278.

Bailey, N.W., Moran, P.A. & Hennig, R.M. (2017) Divergent mechanisms of acoustic mate recognition between closely related field cricket species (Teleogryllus spp.). Animal Behaviour, 130, 17–25.

Bao, W., Kojima, K.K. & Kohany, O. (2015) Repbase Update, a database of repetitive elements in eukaryotic genomes. Mobile Dna, 6, 1–6.

Bay, R.A., Arnegard, M.E., Conte, G.L., Best, J., Bedford, N.L., McCann, S.R., et al. (2017) Genetic coupling of female mate choice with polygenic ecological divergence facilitates stickleback speciation. Current Biology, 27, 3344–3349.

Benson, G. (1999) Tandem repeats finder: a program to analyze DNA sequences. Nucleic acids research, 27, 573–580.

Blankers, T., Oh, K.P. & Shaw, K.L. (2019) Parallel genomic architecture underlies repeated sexual signal divergence in Hawaiian Laupala crickets. Proceedings of the Royal Society B, 286, 20191479.

Bloch, N.I., Corral-López, A., Buechel, S.D., Kotrschal, A., Kolm, N. & Mank, J.E. (2018) Early neurogenomic response associated with variation in guppy female mate preference. Nature ecology & evolution, 2, 1772–1781.

Boeckmann, B., Bairoch, A., Apweiler, R., Blatter, M.-C., Estreicher, A., Gasteiger, E., et al. (2003) The SWISS-PROT protein knowledgebase and its supplement TrEMBL in 2003. Nucleic acids research, 31, 365–370.

Bretman, A., Lawniczak, M.K., Boone, J. & Chapman, T. (2010) A mating plug protein reduces early female remating in Drosophila melanogaster. Journal of insect physiology, 56, 107–113.

Buchfink, B., Reuter, K. & Drost, H.-G. (2021) Sensitive protein alignments at tree-of-life scale using DIAMOND. Nature methods, 18, 366–368.

Cantalapiedra, C.P., Hernández-Plaza, A., Letunic, I., Bork, P. & Huerta-Cepas, J. (2021) eggNOG-mapper v2: functional annotation, orthology assignments, and domain prediction at the metagenomic scale. Molecular biology and evolution, 38, 5825–5829.

Chen, S., Zhou, Y., Chen, Y. & Gu, J. (2018) fastp: an ultra-fast all-in-one FASTQ preprocessor. Bioinformatics, 34, i884–i890.

Cheng, H., Jarvis, E.D., Fedrigo, O., Koepfli, K.-P., Urban, L., Gemmell, N.J., et al. (2022) Haplotype-resolved assembly of diploid genomes without parental data. Nature biotechnology, 40, 1332–1335.

Chowdhury, T., Calhoun, R.M., Bruch, K. & Moehring, A.J. (2020) The fruitless gene affects female receptivity and species isolation. Proceedings of the Royal Society B, 287, 20192765.

Combs, P.A., Krupp, J.J., Khosla, N.M., Bua, D., Petrov, D.A., Levine, J.D., et al. (2018) Tissue-specific cis-regulatory divergence implicates eloF in inhibiting interspecies mating in Drosophila. Current biology, 28, 3969–3975.

Danecek, P., Auton, A., Abecasis, G., Albers, C.A., Banks, E., DePristo, M.A., et al. (2011) The variant call format and VCFtools. Bioinformatics, 27, 2156–2158.

Danecek, P., Bonfield, J.K., Liddle, J., Marshall, J., Ohan, V., Pollard, M.O., et al. (2021) Twelve years of SAMtools and BCFtools. Gigascience, 10, giab008.

Dobin, A., Davis, C.A., Schlesinger, F., Drenkow, J., Zaleski, C., Jha, S., et al. (2013) STAR: ultrafast universal RNA-seq aligner. Bioinformatics, 29, 15–21.

Dolgalev, I. (2022) babelgene: Gene Orthologs for Model Organisms in a Tidy Data Format.

Felsenstein, J. (1981) Skepticism towards Santa Rosalia, or why are there so few kinds of animals? Evolution, 124–138.

Fischer, E.K., Hauber, M.E. & Bell, A.M. (2021) Back to the basics? Transcriptomics offers integrative insights into the role of space, time and the environment for gene expression and behaviour. Biology Letters, 17, 20210293.

Flynn, J.M., Hubley, R., Goubert, C., Rosen, J., Clark, A.G., Feschotte, C., et al. (2020) RepeatModeler2 for automated genomic discovery of transposable element families. Proceedings of the National Academy of Sciences, 117, 9451–9457.

Fraser, H.B. (2022) Existing methods are effective at measuring natural selection on gene expression. Nature Ecology & Evolution, 6, 1836–1837.

Greenacre, M.L., Ritchie, M.G., Byrne, B.C. & Kyriacou, C.P. (1993) Female song preference and the period gene in Drosophila. Behavior genetics, 23, 85–90.

Gross, C., Chang, C.-W., Kelly, S.M., Bhattacharya, A., McBride, S.M., Danielson, S.W., et al. (2015) Increased expression of the PI3K enhancer PIKE mediates deficits in synaptic plasticity and behavior in fragile X syndrome. Cell reports, 11, 727–736.

Guan, D., McCarthy, S.A., Wood, J., Howe, K., Wang, Y. & Durbin, R. (2020) Identifying and removing haplotypic duplication in primary genome assemblies. Bioinformatics, 36, 2896– 2898.

Harrison, P.W., Amode, M.R., Austine-Orimoloye, O., Azov, A.G., Barba, M., Barnes, I., et al. (2024) Ensembl 2024. Nucleic acids research, 52, D891–D899.

Hoy, R.R. (1974) Genetic control of acoustic behavior in crickets. American Zoologist, 14, 1067–1080.

Hoy, R.R., Hahn, J. & Paul, R.C. (1977) Hybrid cricket auditory behavior: evidence for genetic coupling in animal communication. Science, 195, 82–84.

Huerta-Cepas, J., Szklarczyk, D., Heller, D., Hernández-Plaza, A., Forslund, S.K., Cook, H., et al. (2019) eggNOG 5.0: a hierarchical, functionally and phylogenetically annotated orthology resource based on 5090 organisms and 2502 viruses. Nucleic acids research, 47, D309–D314.

Immonen, E. & Ritchie, M.G. (2012) The genomic response to courtship song stimulation in female Drosophila melanogaster. Proceedings of the Royal Society B: Biological Sciences, 279, 1359–1365.

Jacob, P.F. & Hedwig, B. (2019) Structure, Activity and Function of a Singing CPG Interneuron Controlling Cricket Species-Specific Acoustic Signaling. The Journal of Neuroscience, 39, 96–111.

Kanehisa, M., Sato, Y., Kawashima, M., Furumichi, M. & Tanabe, M. (2016) KEGG as a reference resource for gene and protein annotation. Nucleic acids research, 44, D457–D462.

Kataoka, K., Minei, R., Ide, K., Ogura, A., Takeyama, H., Takeda, M., et al. (2020) The draft genome dataset of the Asian cricket Teleogryllus occipitalis for molecular research toward entomophagy. Frontiers in Genetics, 11, 470.

Keagy, J., Hofmann, H.A. & Boughman, J.W. (2024) Mate choice in the brain: species differ in how male traits ‘turn on’gene expression in female brains. Proceedings of the Royal Society B, 291, 20240121.

Kim, D., Paggi, J.M., Park, C., Bennett, C. & Salzberg, S.L. (2019) Graph-based genome alignment and genotyping with HISAT2 and HISAT-genotype. Nature biotechnology, 37, 907–915.

Kirkpatrick, M. (1982) Sexual selection and the evolution of female choice. Evolution, 1–12.

Korunes, K.L. & Samuk, K. (2021) pixy: Unbiased estimation of nucleotide diversity and divergence in the presence of missing data. Molecular ecology resources, 21, 1359–1368.

Kyriacou, C.P. (2002) Single gene mutations in Drosophila: what can they tell us about the evolution of sexual behaviour? Genetics of Mate Choice: From Sexual Selection to Sexual Isolation, 197–203.

Kyriacou, C.P. & Hall, J.C. (1980) Circadian rhythm mutations in Drosophila melanogaster affect short-term fluctuations in the male’s courtship song. Proceedings of the National Academy of Sciences, 77, 6729–6733.

Lande, R. (1981) Models of speciation by sexual selection on polygenic traits. Proceedings of the National Academy of Sciences, 78, 3721–3725.

Lawniczak, M.K. & Begun, D.J. (2004) A genome-wide analysis of courting and mating responses in Drosophila melanogaster females. Genome, 47, 900–910.

Lee, S., Nahm, M., Lee, M., Kwon, M., Kim, E., Zadeh, A.D., et al. (2007) The F-actin-microtubule crosslinker Shot is a platform for Krasavietz-mediated translational regulation of midline axon repulsion.

Love, M.I., Huber, W. & Anders, S. (2014) Moderated estimation of fold change and dispersion for RNA-seq data with DESeq2. Genome Biology, 15, 550.

Lung, O. & Wolfner, M.F. (2001) Identification and characterization of the major Drosophila melanogaster mating plug protein. Insect biochemistry and molecular biology, 31, 543–551.

Lynch, K.S., Ramsey, M.E. & Cummings, M.E. (2012) The mate choice brain: comparing gene profiles between female choice and male coercive poeciliids. Genes, Brain and Behavior, 11, 222–229.

Mack, K.L. & Nachman, M.W. (2017) Gene regulation and speciation. Trends in Genetics, 33, 68–80.

Malinsky, M., Matschiner, M. & Svardal, H. (2021) Dsuite-Fast D-statistics and related admixture evidence from VCF files. Molecular ecology resources, 21, 584–595.

Manni, M., Berkeley, M.R., Seppey, M. & Zdobnov, E.M. (2021) BUSCO: assessing genomic data quality and beyond. Current Protocols, 1, e323.

Martin, S.H., Davey, J.W. & Jiggins, C.D. (2015) Evaluating the use of ABBA–BABA statistics to locate introgressed loci. Molecular biology and evolution, 32, 244–257.

Martinek, S., Inonog, S., Manoukian, A.S. & Young, M.W. (2001) A role for the segment polarity gene shaggy/GSK-3 in the Drosophila circadian clock. Cell, 105, 769–779.

Mendelson, T.C. & Shaw, K.L. (2005) Rapid speciation in an arthropod. Nature, 433, 375– 376.

Moran, P.A., Hunt, J., Mitchell, C., Ritchie, M.G. & Bailey, N.W. (2019) Behavioural mechanisms of sexual isolation involving multiple modalities and their inheritance. Journal of evolutionary Biology, 32, 243–258.

Moran, P.A., Pascoal, S., Cezard, T., Risse, J.E., Ritchie, M.G. & Bailey, N.W. (2018) Opposing patterns of intraspecific and interspecific differentiation in sex chromosomes and autosomes. Molecular Ecology, 27, 3905–3924.

Moran, P.A., Ritchie, M.G. & Bailey, N.W. (2017) A rare exception to Haldane’s rule: Are X chromosomes key to hybrid incompatibilities? Heredity, 118, 554–562.

Moriya, Y., Itoh, M., Okuda, S., Yoshizawa, A.C. & Kanehisa, M. (2007) KAAS: an automatic genome annotation and pathway reconstruction server. Nucleic acids research, 35, W182– W185.

Muralidhar, M. & Thomas, J.B. (1993) The Drosophila bendless gene encodes a neural protein related to ubiquitin-conjugating enzymes. Neuron, 11, 253–266.

Oh, C.E., McMahon, R., Benzer, S. & Tanouye, M.A. (1994) bendless, a Drosophila gene affecting neuronal connectivity, encodes a ubiquitin-conjugating enzyme homolog. Journal of Neuroscience, 14, 3166–3179.

Parker, D.J., Gardiner, A., Neville, M.C., Ritchie, M.G. & Goodwin, S.F. (2014) The evolution of novelty in conserved genes; evidence of positive selection in the Drosophila fruitless gene is localised to alternatively spliced exons. Heredity, 112, 300–306.

Pascoal, S., Liu, X., Fang, Y., Paterson, S., Ritchie, M.G., Rockliffe, N., et al. (2018) Increased socially mediated plasticity in gene expression accompanies rapid adaptive evolution. Ecology Letters, 21, 546–556.

Pascoal, S., Risse, J.E., Zhang, X., Blaxter, M., Cezard, T., Challis, R.J., et al. (2020) Field cricket genome reveals the footprint of recent, abrupt adaptation in the wild. Evolution Letters, 4, 19–33.

Paysan-Lafosse, T., Blum, M., Chuguransky, S., Grego, T., Pinto, B.L., Salazar, G.A., et al. (2023) InterPro in 2022. Nucleic acids research, 51, D418–D427.

Picard toolkit. (2018) Broad Institute, GitHub repository.

Quinlan, A.R. & Hall, I.M. (2010) BEDTools: a flexible suite of utilities for comparing genomic features. Bioinformatics, 26, 841–842.

Ramsey, M.E., Maginnis, T.L., Wong, R.Y., Brock, C. & Cummings, M.E. (2012) Identifying Context-Specific Gene Profiles of Social, Reproductive, and Mate Preference Behavior in a Fish Species with Female Mate Choice. Frontiers in Neuroscience, 6.

Rhie, A., Walenz, B.P., Koren, S. & Phillippy, A.M. (2020) Merqury: reference-free quality, completeness, and phasing assessment for genome assemblies. Genome biology, 21, 1–27.

Rideout, E.J., Billeter, J.-C. & Goodwin, S.F. (2007) The sex-determination genes fruitless and doublesex specify a neural substrate required for courtship song. Current Biology, 17, 1473– 1478.

Ritchie, M.G. & Butlin, R.K. (2024) Genetic Coupling of Mate Recognition Systems in the Genomic Era. Cold Spring Harbor Perspectives in Biology, a041437.

Rossi, M., Hausmann, A.E., Alcami, P., Moest, M., Roussou, R., Van Belleghem, S.M., et al. (2024) Adaptive introgression of a visual preference gene. Science, 383, 1368–1373.

Rossi, M., Hausmann, A.E., Thurman, T.J., Montgomery, S.H., Papa, R., Jiggins, C.D., et al. (2020) Visual mate preference evolution during butterfly speciation is linked to neural processing genes. Nature Communications, 11, 4763.

Sabatier, L., Jouanguy, E., Dostert, C., Zachary, D., Dimarcq, J.-L., Bulet, P., et al. (2003) Pherokine-2 and-3: Two Drosophila molecules related to pheromone/odor-binding proteins induced by viral and bacterial infections. European journal of biochemistry, 270, 3398–3407.

Sakisaka, T., Yamamoto, Y., Mochida, S., Nakamura, M., Nishikawa, K., Ishizaki, H., et al. (2008) Dual inhibition of SNARE complex formation by tomosyn ensures controlled neurotransmitter release. The Journal of cell biology, 183, 323–337.

Sánchez-Ramírez, S. & Cutter, A. (2021) CompMap: an allele-specific expression read-counter based on competitive mapping. bioRxiv, 2021–02.

Sánchez-Ramírez, S., Weiss, J.G., Thomas, C.G. & Cutter, A.D. (2021) Widespread misregulation of inter-species hybrid transcriptomes due to sex-specific and sex-chromosome regulatory evolution. PLoS genetics, 17, e1009409.

Sayers, E.W., Bolton, E.E., Brister, J.R., Canese, K., Chan, J., Comeau, D.C., et al. (2021) Database resources of the national center for biotechnology information. Nucleic Acids Research, 50, D20–D26.

Sedlazeck, F.J., Rescheneder, P., Smolka, M., Fang, H., Nattestad, M., Von Haeseler, A., et al. (2018) Accurate detection of complex structural variations using single-molecule sequencing. Nature methods, 15, 461–468.

Shaw, K.L., Cooney, C.R., Mendelson, T.C., Ritchie, M.G., Roberts, N.S. & Yusuf, L.H. (2024) How Important Is Sexual Isolation to Speciation? Cold Spring Harbor Perspectives in Biology, a041427.

Shaw, K.L. & Lesnick, S.C. (2009) Genomic linkage of male song and female acoustic preference QTL underlying a rapid species radiation. Proceedings of the National Academy of Sciences, 106, 9737–9742.

Shumate, A. & Salzberg, S.L. (2021) Liftoff: accurate mapping of gene annotations. Bioinformatics, 37, 1639–1643.

Subkhangulova, A., Gonzalez-Lozano, M.A., Groffen, A.J., Weering, J.R. van, Smit, A.B., Toonen, R.F., et al. (2023) Tomosyn affects dense core vesicle composition but not exocytosis in mammalian neurons. Elife, 12, e85561.

Tan, Y., Yu, D., Busto, G.U., Wilson, C. & Davis, R.L. (2013) Wnt signaling is required for long-term memory formation. Cell reports, 4, 1082–1089.

Tarailo-Graovac, M. & Chen, N. (2009) Using RepeatMasker to identify repetitive elements in genomic sequences Curr Protoc Bioinformatics.

Tarasov, A., Vilella, A.J., Cuppen, E., Nijman, I.J. & Prins, P. (2015) Sambamba: fast processing of NGS alignment formats. Bioinformatics, 31, 2032–2034.

Tillich, M., Lehwark, P., Pellizzer, T., Ulbricht-Jones, E.S., Fischer, A., Bock, R., et al. (2017) GeSeq–versatile and accurate annotation of organelle genomes. Nucleic acids research, 45, W6–W11.

Uliano-Silva, M., Ferreira, J.G.R., Krasheninnikova, K., Formenti, G., Abueg, L., Torrance, J., et al. (2023) MitoHiFi: a python pipeline for mitochondrial genome assembly from PacBio high fidelity reads. BMC bioinformatics, 24, 288.

Unbehend, M., Kozak, G.M., Koutroumpa, F., Coates, B.S., Dekker, T., Groot, A.T., et al. (2021) bric à brac controls sex pheromone choice by male European corn borer moths. Nature Communications, 12, 2818.

Uthaman, S.B., Godenschwege, T.A. & Murphey, R.K. (2008) A mechanism distinct from highwire for the Drosophila ubiquitin conjugase bendless in synaptic growth and maturation. Journal of Neuroscience, 28, 8615–8623.

Vasimuddin, M., Misra, S., Li, H. & Aluru, S. (2019) Efficient architecture-aware acceleration of BWA-MEM for multicore systems. In 2019 IEEE international parallel and distributed processing symposium (IPDPS). IEEE, pp. 314–324.

Wehrli, M., Dougan, S.T., Caldwell, K., O’Keefe, L., Schwartz, S., Vaizel-Ohayon, D., et al. (2000) arrow encodes an LDL-receptor-related protein essential for Wingless signalling. Nature, 407, 527–530.

Wellenreuther, M. & Sánchez-Guillén, R.A. (2016) Nonadaptive radiation in damselflies. Evolutionary Applications, 9, 103–118.

Wolf, F.W., Eddison, M., Lee, S., Cho, W. & Heberlein, U. (2007) GSK-3/Shaggy regulates olfactory habituation in Drosophila. Proceedings of the National Academy of Sciences, 104, 4653–4657.

Woolley, S.C. & Doupe, A.J. (2008) Social context–induced song variation affects female behavior and gene expression. PLoS biology, 6, e62.

Wu, M., Walser, J.-C., Sun, L. & Kölliker, M. (2020) The genetic mechanism of selfishness and altruism in parent-offspring coadaptation. Science advances, 6, eaaw0070.

Yamamoto, Y., Mochida, S., Miyazaki, N., Kawai, K., Fujikura, K., Kurooka, T., et al. (2010) Tomosyn inhibits synaptotagmin-1-mediated step of Ca2+-dependent neurotransmitter release through its N-terminal WD40 repeats. Journal of Biological Chemistry, 285, 40943–40955.

Ye, W., Dang, J., Xie, L. & Huang, Y. (2008) Complete mitochondrial genome of Teleogryllus emma (Orthoptera: Gryllidae) with a new gene order in Orthoptera.

Yizhar, O., Matti, U., Melamed, R., Hagalili, Y., Bruns, D., Rettig, J., et al. (2004) Tomosyn inhibits priming of large dense-core vesicles in a calcium-dependent manner. Proceedings of the National Academy of Sciences, 101, 2578–2583.

Ylla, G., Nakamura, T., Itoh, T., Kajitani, R., Toyoda, A., Tomonari, S., et al. (2021) Insights into the genomic evolution of insects from cricket genomes. Communications biology, 4, 733.

York, R.A., Patil, C., Abdilleh, K., Johnson, Z.V., Conte, M.A., Genner, M.J., et al. (2018) Behavior-dependent cis regulation reveals genes and pathways associated with bower building in cichlid fishes. Proceedings of the National Academy of Sciences, 115, E11081–E11090.

Yu, G., Wang, L., Han, Y. & He clusterProfiler, Q. (2011) An R package for comparing biological themes among gene clusters., 2012, 16. *DOI:* 10.1089/omi, 284–287.

Yuan, Q., Lin, F., Zheng, X. & Sehgal, A. (2005) Serotonin modulates circadian entrainment in Drosophila. Neuron, 47, 115–127.

Yusuf, L.H., Pascoal, S., Moran, P.A. & Bailey, N.W. (2024) Testing the genomic overlap between intraspecific mating traits and interspecific mating barriers. Evolution Letters, qrae042.

Zhang, X., Blaxter, M., Wood, J.M., Tracey, A., McCarthy, S., Thorpe, P., et al. (2024) Temporal genomics in Hawaiian crickets reveals compensatory intragenomic coadaptation during adaptive evolution. Nature Communications, 15, 5001.

Zhang, X., Rayner, J.G., Blaxter, M. & Bailey, N.W. (2021) Rapid parallel adaptation despite gene flow in silent crickets. Nature Communications, 12, 50.

Zhou, C., McCarthy, S.A. & Durbin, R. (2023) YaHS: yet another Hi-C scaffolding tool. Bioinformatics, 39, btac808.

Zhu, A., Ibrahim, J.G. & Love, M.I. (2019) Heavy-tailed prior distributions for sequence count data: removing the noise and preserving large differences. Bioinformatics, 35, 2084–2092.

